# Cdc42 regulates cytokine expression and trafficking in bronchial epithelial cells

**DOI:** 10.1101/2022.11.09.515863

**Authors:** Rowayna Shouib, Gary Eitzen

## Abstract

Airway epithelial cells can respond to incoming pathogens, allergens and stimulants through the secretion of cytokines and chemokines. These pro-inflammatory mediators activate inflammatory signaling cascades that allow a robust immune response to be mounted. However, uncontrolled production and release of cytokines and chemokines can result in chronic inflammation and appears to be an underlying mechanism for the pathogenesis of pulmonary disorders such as asthma and COPD. The Rho GTPase, Cdc42, is an important signaling molecule that we hypothesize can regulate cytokine production and release from epithelial cells. We treated BEAS-2B lung epithelial cells with a set of stimulants to activate inflammatory pathways and cytokine release. The production, trafficking and secretion of cytokines were assessed when Cdc42 was pharmacologically inhibited with ML141 drug or silenced with lentiviral-mediated shRNA knockdown. We found that Cdc42 inhibition with ML141 differentially affected gene expression of a subset of cytokines; transcription of IL-6 and IL-8 were increased while MCP-1 was decreased. However, Cdc42 inhibition or depletion disrupted IL-8 trafficking and reduced its secretion even though transcription was increased. Cytokines transiting through the Golgi were particularly affected by Cdc42 disruption. Our results define a role for Cdc42 in the regulation of cytokine production and release in airway epithelial cells. This underscores the role of Cdc42 in coupling receptor activation to downstream gene expression and also as a regulator of cytokine secretory pathways.

**Short Summary:** Cytokine secretion from airway epithelial cells contributes to the pathogenesis of disease. We show that Cdc42 regulates cytokine gene expression and is required for cytokine secretion via control of transport through the Golgi complex.

## INTRODUCTION

Airway epithelial cells (AECs) that line the pulmonary tracts provide a relatively impermeable physical barrier and maintain the conduit for airflow to and from the alveoli. In addition to barrier function, AECs play an important role in immunity. Bronchial epithelial cells in particular are involved in immunological processes in the lungs, providing protection against inhaled pathogens through engaging host immune defense mechanisms (1). This process is orchestrated through the upregulation and release of pro-inflammatory cytokines and chemokines, which signal downstream recruitment and activation of immune cells. AECs are also involved in the dysfunctional regulation of inflammatory processes in the airways associated with major airway disorders such as asthma and chronic obstructive pulmonary disorder (COPD). For instance, epithelial-derived cytokines and chemokines such as IL-25, IL-33, GM-CSF and TSLP, are produced upon exposure to allergens and contribute to the state of allergic inflammation and airway damage in asthma (2). The bronchial epithelium of asthmatics also displays increased expression of pro-inflammatory transcription factors such as NFκB and AP-1 (3).

Bronchial epithelial cells are equipped with numerous cell surface receptors that enable recognition of pathogens, allergens and pollutants and the activation of pro-inflammatory pathways. These receptors include Toll-like receptors (TLR), protease-activated receptors (PAR) and other G-protein coupled receptors (GPCR) (4). TLRs are a form of pattern-recognition receptors (PRRs) which respond to conserved structures known as pathogen-associated molecular patterns (PAMPs) derived from bacterial, viral, fungal or parasitic sources (5). Upon binding of the PAMPs to their corresponding TLRs, a signaling cascade is initiated that ultimately results in release of pro-inflammatory mediators. G-protein coupled receptors such as PAR-2 are activated by serine proteases present in household allergens such as cockroach extract allergen which cleaves the N-terminal exodomain of the PAR-2 receptor activating the respective signaling cascade that leads to inflammation (6). Therefore, exposure to this cockroach allergen is known to be associated with the onset and progression of asthma especially amongst young children (7). Another key surface receptor on epithelial cells is the receptor for the TNF-α cytokine. TNF-α binds TNFR1, which is ubiquitously expressed and mediates the majority of the downstream effects of TNF-α, and/or TNFR2, which is expressed exclusively in lymphocytes and endothelial cells (8). Upon binding to TNFR1, a pathway is activated that results in NF-κB activation and nuclear translocation and the transcription of additional pro-inflammatory cytokines and chemokines.

It is clear that the process of transcription and secretion of pro-inflammatory mediators allows AECs to modulate early inflammatory events, thereby initiating inflammation or exacerbating existing inflammation if unregulated (2). However, the exact mechanism by which the engagement of epithelial cell receptors with pathogens and allergic stimuli is coupled to the release of pro-inflammatory mediators remains largely unexplained. Rho proteins, small GTPases of the Rho family, can act as molecular switches within the cell and are known to be involved in cytoskeletal remodeling, vesicle trafficking and signal transduction (9). These proteins exist within cells in either the active GTP-bound form and the inactive GDP-bound form and are proposed to be regulators in inflammatory signaling pathways and gene expression. For instance, Cdc42 has been shown to be required for pro-inflammatory gene expression in human senescent endothelial cells (10). Research has pointed to the involvement of Rho proteins in the pathogenesis of asthma. In an animal model of asthma, allergen-sensitized mice treated with the Rac1 inhibitor, NSC23766, showed reduced bronchial inflammation compared to untreated mice (11). Although Cdc42 has also been proposed to be involved in the secretory trafficking and exocytosis process in a variety of conditions, its role in affecting the trafficking and secretion of pro-inflammatory cytokines and chemokines in lung epithelial cells remains unexamined. However, a role for Cdc42 in secretion has been demonstrated in the release of CCL2 chemokine from PNMECs (pulmonary neuroendocrine cells) (12) and IFN-γ cytokine from T-cells at the immunological synapse through mediating actin depolymerization (13). Therefore, Rho GTPases, and Cdc42 in particular, are interesting candidates in the regulation of cytokine and chemokine upregulation and secretion in bronchial epithelial cells.

BEAS-2B cells are an immortalized human bronchial epithelial cell line that has been widely used as an in vitro cell model for studies related to respiratory disease (14). In this study, we induced a state of inflammation in BEAS-2B cells through the treatment with different allergens and stimulants such as cockroach extract, polyinosinic-polycytidylic acid (poly(I:C)), and TNF-α. The state of inflammation was assessed through quantifying the production and release of pro-inflammatory cytokines from stimulated cells. Subsequently, we examined the role of Rho proteins in the pro-inflammatory pathways through the use of selective small molecule inhibitors. We found that Cdc42 was involved in the regulation of cytokines, in both the production and trafficking in lung epithelial cells. Our results show that the Rho protein, Cdc42, plays a pivotal role in the signal transduction events that lead to upregulation and increased release of cytokines from lung epithelial cells.

## RESULTS

### Various stimuli activate cytokine production and release in BEAS-2B cells

Bronchial epithelial cells provide an immunological barrier in the airways and can respond to a variety of noxious substances through the release of pro-inflammatory cytokines (2, 3). We examined the response of the human bronchial epithelial cell line, BEAS-2B, to three stimuli that activate different receptors: poly(I:C), which activates TLR3; cockroach extract, which activates the GPCR PAR1; and TNF-α, which activates TNFR. Extracellular supernatants were collected after 8 h of stimulation with these ligands and analyzed for 15 cytokines using a commercially sourced multiplex assay (**Table S1**, *supplementary material*). Of the fifteen factors, IL-8, MCP-1 and IL-6 gave reproducibly detectable responses, with TNF-α eliciting the most significant increase in released cytokines (**Figure 1A**). Examination of cytokine gene expression by qPCR showed that stimulation with poly(I:C) and cockroach extract induced a 3-fold and 2-fold increase in IL-8 cytokine gene expression, respectively, while TNF-α increased IL-8 mRNA > 500-fold (**Figure 1B**). These results show that TNF-α is a potent activator of cytokine production and release in BEAS-2B lung epithelial cells.

**Figure 1.**
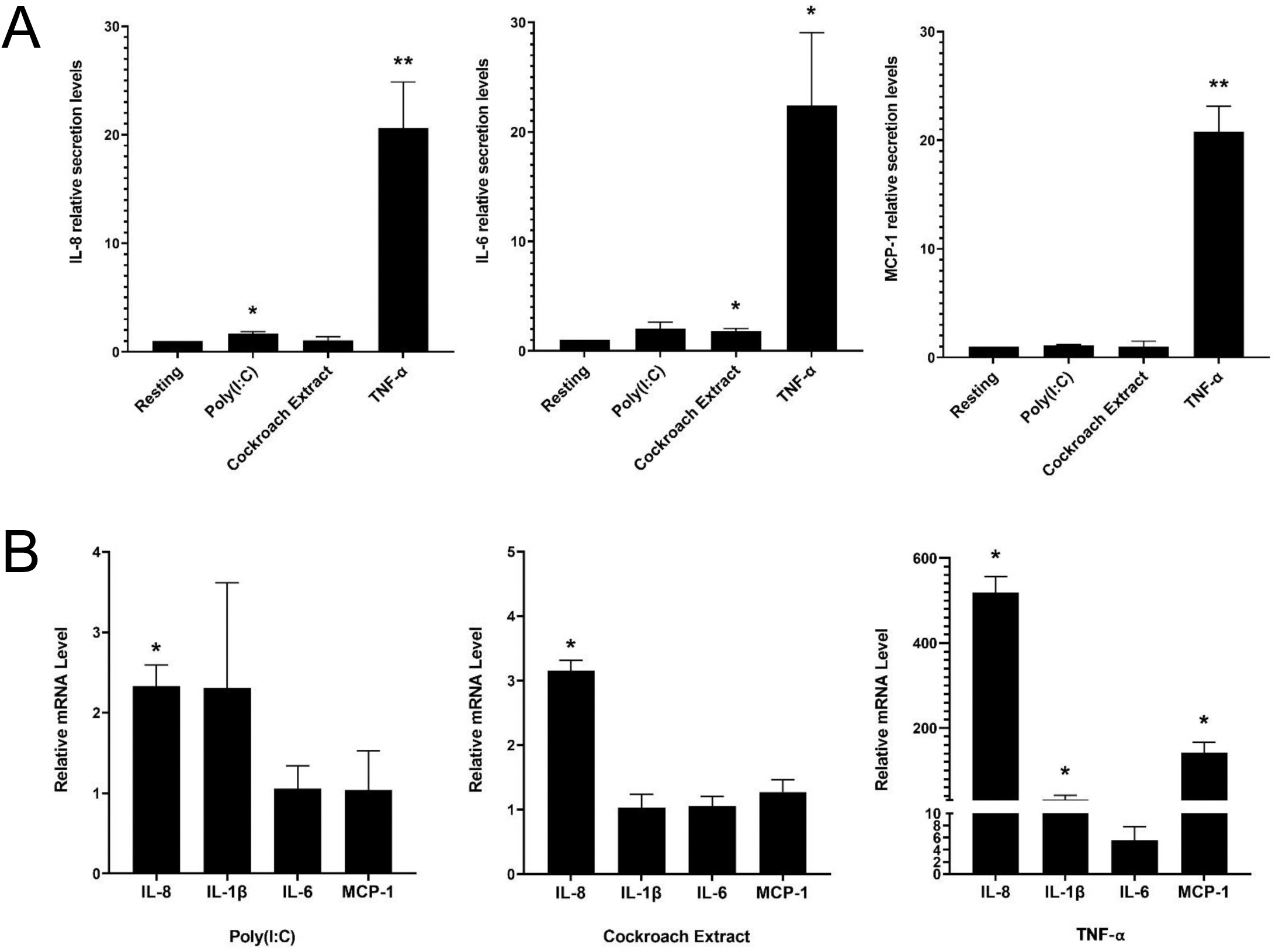
BEAS-2B cells respond to pro-inflammatory stimuli with upregulated cytokine production and release. BEAS-2B cells were stimulated with 10 μg/mL poly(I:C), 20 μg/mL cockroach extract and 10 ng/mL TNF-α. **A)** Relative levels of IL-8, IL-1β, IL-6 and MCP-1 in cell culture media after 8 h of stimulation as detected by multiplex assay. Values represent secretion levels normalized to unstimulated samples. **B)** Fold change in mRNA levels of cytokine genes after 4 h of stimulation as detected by qPCR. Values represent ΔΔCt fold changes normalized to unstimulated samples. Bars are the mean ± s.e.m.; n=3; ** p <0.01, * p < 0.05 by unpaired two-tailed Student’s t-test.

Morphological changes to BEAS-2B cells after stimulation with the three agonists were examined by immunofluorescence microscopy. Cells were stimulated for 4 hr, fixed and stained for F-actin and microtubules to show the cell structure, and IL-8 to show cytokine expression and trafficking (**Figure 2A**). The imaging showed an overall increase in IL-8 staining upon stimulation with all three agonists compared to the resting (unstimulated) control cells. There was no change in microtubule structures, while an increase in perinuclear F-actin puncta was observed after stimulation, especially with cockroach extract and TNF-α (**Figure 2A**, *arrows*). In particular, TNF-α induced a prominent increase in Golgi and post-Golgi cytokine staining. In addition, the staining pattern of NF-κB, a prominent cytokine transcription factor, was assessed. Prior to stimulation, the NF-κB p65 subunit was exclusively cytosolic, however, upon cell stimulation by agonists it was translocated to the nucleus. Although cells stimulated with cockroach extract and poly(I:C) displayed only minimal NF-κB staining in the nucleus, a robust translocation was observed following TNF-α stimulation (**Figure 2B** and **2C**). Therefore, all further analyses were performed using TNF-α as the stimulus.

**Figure 2.**
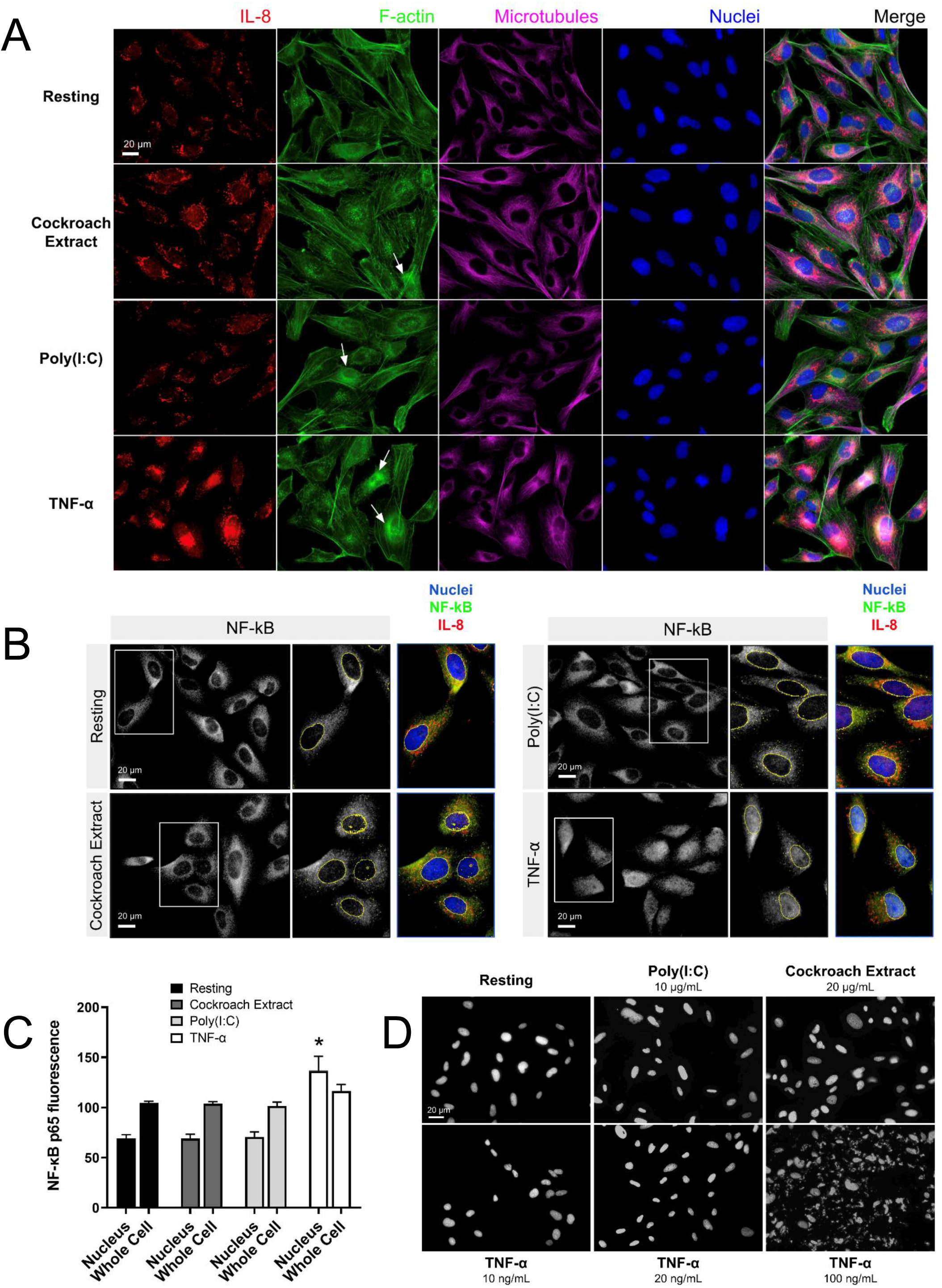
Immunofluorescence microscopy shows cytokine upregulation in response to treatment with pro-inflammatory stimuli. **A and B)** BEAS-2B cells were serum-starved for 16 h, then stimulated with 20 μg/mL cockroach extract, 10 μg/mL poly(I:C) or 10 ng/mL TNF-α. **A)** Cells were stimulated for 4 h then fixed and stained with anti-IL8 and anti-β–tubulin antibodies, Alexa546-phalloidin (F-actin), and DAPI (nuclei). **B and C)** Quantification of NF-κB levels in the nucleus. Cells were stimulated for 30 min then fixed and stained with anti-NF-κB antibodies and DAPI (nuclei) for immunofluorescence (*panel B*) and levels of NF-κB staining in the nuclei or whole cells (nuclei and cytosolic regions) was quantified (*panel C*). Bars are the mean ± s.e.m.; n=4; * p < 0.05 by unpaired two-tailed Student’s t-test. **D)** DAPI staining of BEAS-2B cells to assess apoptosis. Cells were treated with the indicated stimuli for 4 h then fixed and stained with DAPI to analyze nuclear fragmentation which indicates apoptosis.

TNF-α is well known to activate cell death by apoptosis in addition to the upregulation of cytokine production via the NF-κB pathway (15). To determine whether cell death was activated in BEAS-2B cells, we examined nuclear fragmentation after TNF-α treatment. Cells treated with 10 ng/mL TNF-α (commonly used in this study to stimulate cytokine production) showed a minimal number of apoptotic cells, comparable to the untreated control cells (**Figure 2D**). Significant apoptosis of BEAS-2B cells was observed only at high concentrations of TNF-α (100 ng/mL). Since there was a significant increase in nuclear-localized NF-κB upon TNF-α stimulation (**Figure 2B** and **2C**), this suggests that BEAS-2B cells predominantly respond to TNF-α with cell survival and upregulation of NF-κB-directed transcription.

### Cdc42 regulates lung epithelial cell cytokine production and release

Next, we wanted to determine whether Rho GTPases affect cytokine production and release from lung epithelial cells. Several studies have shown links between the Rho GTPases and NF-κB-regulated cytokine production, but these have shown both positive or negative regulatory roles, depending on the context (16–18, *reviewed in* 19). We examined the role of Rho GTPases using specific Rho inhibitors. BEAS-2B cells were pretreated with Rhosin, EHT1864 and ML141 to inhibit RhoA, Rac1 and Cdc42, respectively (20–22), then stimulated with 10 ng/mL TNF-α for 4 hr to elicit a response. mRNA was isolated and transcript levels were examined by qPCR. RhoA and Rac1 inhibitors induced consistent, though small, increases in cytokine gene expression. The Cdc42 inhibitor, ML141, significantly increased in IL-6, IL-8 and IL-1β mRNA levels, but reduced MCP-1 mRNA levels (**Figure 3A**). We also examined the effects of drugs that regulate both cytokine transcription and trafficking, for comparison. The NF-κB inhibitor, BAY 11-7082, significantly reduced of all cytokine transcription while the Golgi trafficking inhibitor, monensin, had minimal effect (**Figure 3A**). This shows that the ML141 effect is not a general drug effect. The effects of ML141 on cytokine secretion were also examined. While IL-8 and IL-6 transcription greatly increased when treated with Cdc42 inhibitor, their secretion was not concomitantly increased (**Figure 3B**). However, MCP-1, which showed a modest decrease in transcription, showed a drastic reduction in secretion when treated with ML141 (**Figure 3B**). ML141 did not affect the translocation of NF-κB into the nucleus, nor the levels of nuclear localized NF-κB compared to vehicle-treated cells (**Figure 3C and 3D**). These results suggest that Cdc42 may negatively regulate transcriptional activation of a subset of cytokines, however, this is not mediated NF-κB pathway. Cdc42 also seems to be required for general cytokine secretion and thus have dual roles in regulating cytokines.

**Figure 3.**
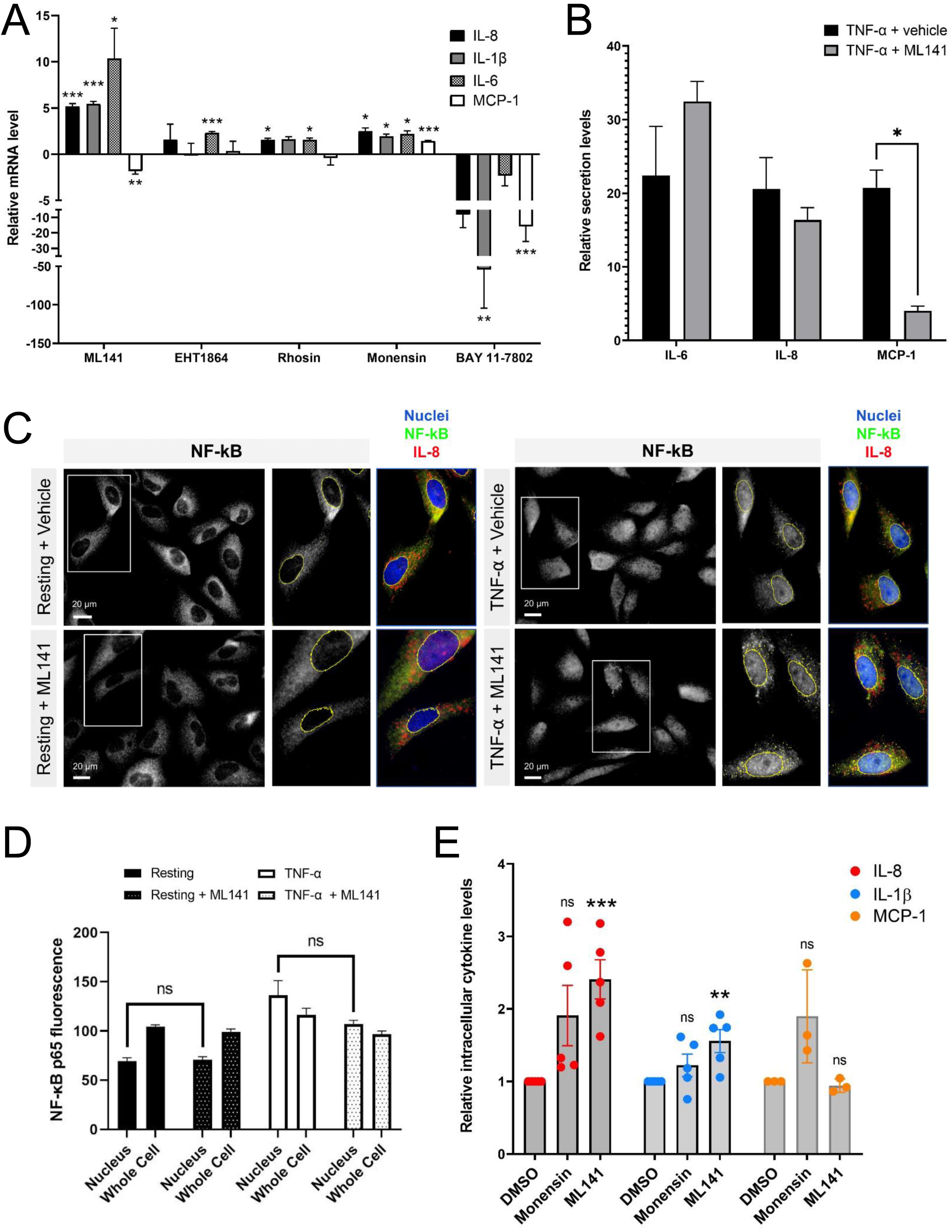
The Cdc42 inhibitor, ML141, affects cytokine production and secretion. **A)** Change in cytokine mRNA levels due to incubation with Rho drugs. BEAS-2B cells were serum-starved for 16 h and pretreated with drugs for 1 h, then stimulated with 10 ng/mL TNF-α for 4 h. Values represent ΔΔCt fold changes in mRNA normalized to vehicle-treated control. Drug concentrations: 20 μM ML141, 10 μM EHT1864, 10 μM rhosin, 2 μM monensin, 10 μM BAY 11-7082. **B)** Relative levels of secreted IL-6, IL-8 and MCP-1 in cell culture media of BEAS-2B cells. Conditioned media was collected after 8 h of stimulation with 10 ng/mL TNF-α +/− 20 μM ML141, and cytokines detected by human cytokine multiplex assay (EveTechnologies™). **C and D)** Quantification of NF-κB levels in the nucleus. BEAS-2B cells were pre-treated with 20 μM ML141 or vehicle (DMSO) for 1 h, then stimulated for 30 min with 10 ng/mL TNF-α. Cells were fixed and stained with NF-κB antibodies for immunofluorescence (*panel C*) and levels of NF-κB staining in the nuclei or whole cells (nuclei and cytosolic regions) was quantified (*panel D*).**E)** Intracellular levels of cytokines in stimulated BEAS-2B cells detected by flow cytometry. Cells were pre-treated with 2 μM monensin, 20 μM ML141 or vehicle (*DMSO*) for 1 h then stimulated with 10 ng/mL TNF-α for 4 h. Cells were fixed and stained with IL-8, IL-1β, and MCP-1 antibodies. Bars are the mean ± s.e.m; n=4 (15 – 20 cells from at least 3 different images); *** p < 0.001, ** p < 0.01, * p < 0.05 by unpaired two-tailed Student’s t-test.

To support these results, we used flow cytometry to quantify intracellular cytokine levels. We used the Golgi transport inhibitor, monensin, as a control to increase intracellular cytokine levels by blocking secretion. As expected, monensin increased intracellular IL-8 and MCP-1 cytokine levels but did not affect IL-1β levels, a cytokine that is constitutively secreted and does not traffic through the Golgi (23). The average level of IL-8 intracellular staining was much higher upon monensin treatment (**Figure 3E**), however variability resulted in this effect not being statistically significant. Interestingly, we found that ML141 treatment resulted in an increase in intracellular levels of both IL-8 and IL-1β (**Figure 3E**), which is concomitant with their upregulated expression (*see Figure 3A*). The increase in IL-1β levels are less likely to be affected by Cdc42 at the Golgi; therefore, the effect observed is likely mediated through gene upregulation. However, the increase in intracellular levels of IL-8 could be a combined result of both increased cytokine transcription and decreased secretion from the cell. We also found that ML141 does not affect MCP-1 intracellular levels (**Figure 3E**) despite reducing MCP-1 gene expression (*see Figure 3A*). This suggests that MCP-1 secretion could be blocked resulting in greater intracellular accumulation.

### Cdc42 regulates lung epithelial cell cytokine trafficking

To further differentiate between Cdc42 effects on transcription vs secretion of cytokines, we performed immunofluorescence microscopy to visualize cytokine trafficking. BEAS-2B cells were either left unstimulated, or stimulated for 4 hours with TNF-α. IL-8 and IL-1β staining increased after TNF-α stimulation; however, their distribution within the cell was distinct. IL-8 showed perinuclear staining and an increase in peripheral tubule reticular staining after stimulation (**Figure 4A**, *upper panels*), while IL-1β showed a general punctate staining pattern (**Figure 4B**, *upper panels*). IL-8 and IL-1β staining did not overlap (**Figure S1**, *supplementary material*). Inhibition of Cdc42 with ML141 resulted in the loss of IL-8 tubule staining, and instead, intense punctate staining was observed (**Figure 4A**, *lower panels*), while the IL-1β staining pattern was essentially not affected (**Figure 4B**, *lower panels*). Tubule staining of IL-8 also occurred when other stimuli were used and these were similarly disrupted by ML141 (**Figure 4A**, *Cockroach Extract, Poly(I:C)*)

**Figure 4.**
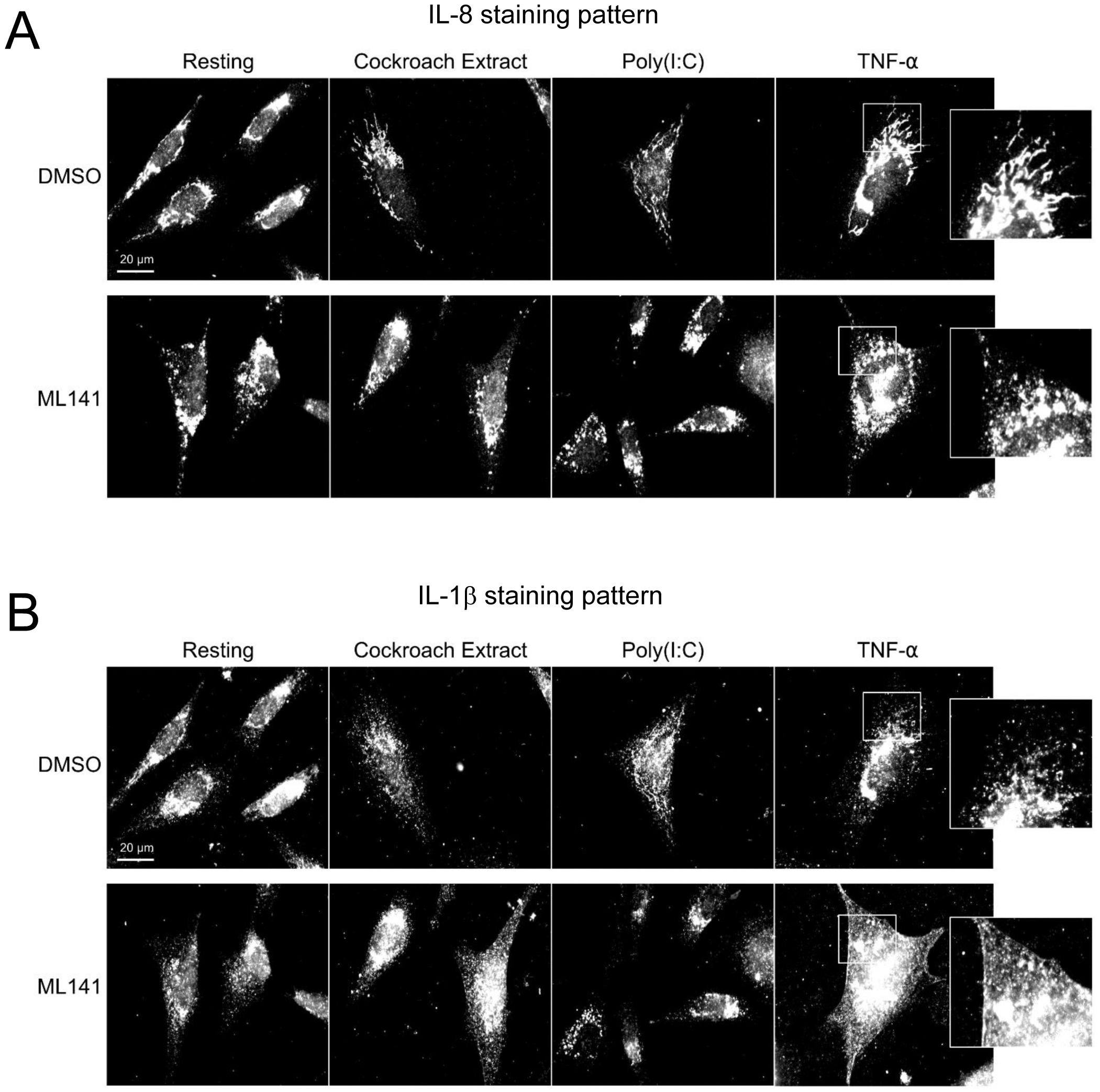
Cdc42 inhibition disrupts cytokine trafficking. BEAS-2B cells were pretreated with 20 μM ML141 or vehicle (DMSO) for 1 h, then stimulated with 10 ng/mL TNF-α for 4 h. Cells were then fixed and stained with cytokine antibodies. **A)** IL-8 staining pattern. **B)** IL-1β staining pattern. Zoomed panels show the two cytokines display different characteristic staining patterns upon stimulation. ML141 affects the IL-8 staining pattern but does not affect IL-1β staining.

We next examined the cellular distribution of cytokines with respect to the Golgi apparatus since their trafficking and secretion is likely to be Golgi-dependent. In addition to IL-8, MCP-1 distribution was examined since these cytokines are differentially affected by ML141. Both IL-8 and MCP-1 showed an increase in Golgi and post-Golgi staining after 4 hours of TNF-α stimulation (**Figure 5A** and **6A**). IL-8 labeled tubules were observed that projected toward the cell periphery (**Figure 5A**, *arrow*), but tubule labeling was not observed for MCP-1 (**Figure 6A**). This suggests that the post-Golgi trafficking pathways of IL-8 and MCP-1 are different. In the presence of monensin, both IL-8 and MCP-1 showed an increase in Golgi staining and reduced post-Golgi staining (**Figure 5B** and **6B**, *monensin*), which defines the Golgi as central to the secretion of both cytokines. Notably, treatment with the Cdc42 inhibitor, ML141, resulted in the loss of post-Golgi tubule staining of IL-8 and increased staining of puncta around the Golgi (**Figure 5B**, *ML141*). Treatment with ML141 resulted in the partial dispersal of the Golgi staining of MCP-1 observed upon treatment with TNF-α, and increased punctate staining pattern around the Golgi (**Figure 6B**, *ML141*). The Golgi marker, GM130, also showed an increase in dispersed punctate staining after ML141 treatment (**Figure 5B** and **6B**, *green channel*). This suggests that Cdc42 is needed for efficient trafficking of cytokines through the Golgi due to its requirement to maintain Golgi structure. A degree of Golgi disruption (apparent from GM130 staining) was also observed in monensin-treated cells, consistent with findings reporting GM130 degradation and Golgi dispersal in response to monensin (24). IL-1β showed a general cytosolic staining pattern since it does not traffic through the Golgi and this was not affected by ML141 treatment (*see Figure 4*).

**Figure 5.**
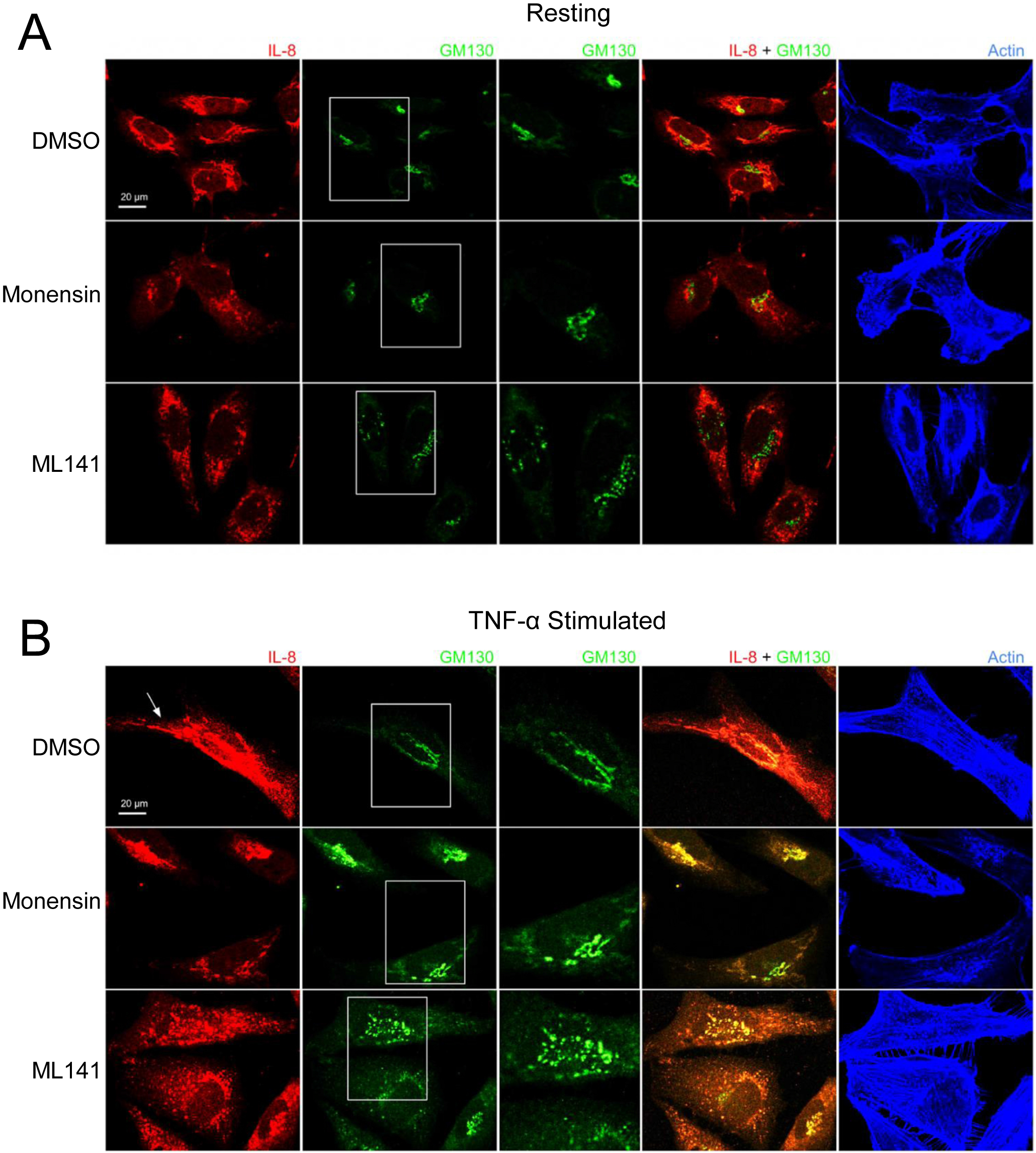
Cdc42 inhibition induces changes in Golgi structure and in IL-8 trafficking. BEAS-2B cells were pre-treated with 20 μM ML141, 2 μM monensin or vehicle (DMSO) for 1 h. **A)** Cells left unstimulated (*resting*) for 4 h. **B)** Cells were stimulated with 10 ng/mL TNF-α for 4 h. Cells were then fixed and stained with IL-8 and GM130 (Golgi marker) antibodies while actin was labeled with phalloidin-iFluor 405 dye. Monensin induced a Golgi-localized staining pattern for IL-8 while ML141 treatment resulted in less Golgi-localized IL-8 and dispersed GM130 staining.

**Figure 6.**
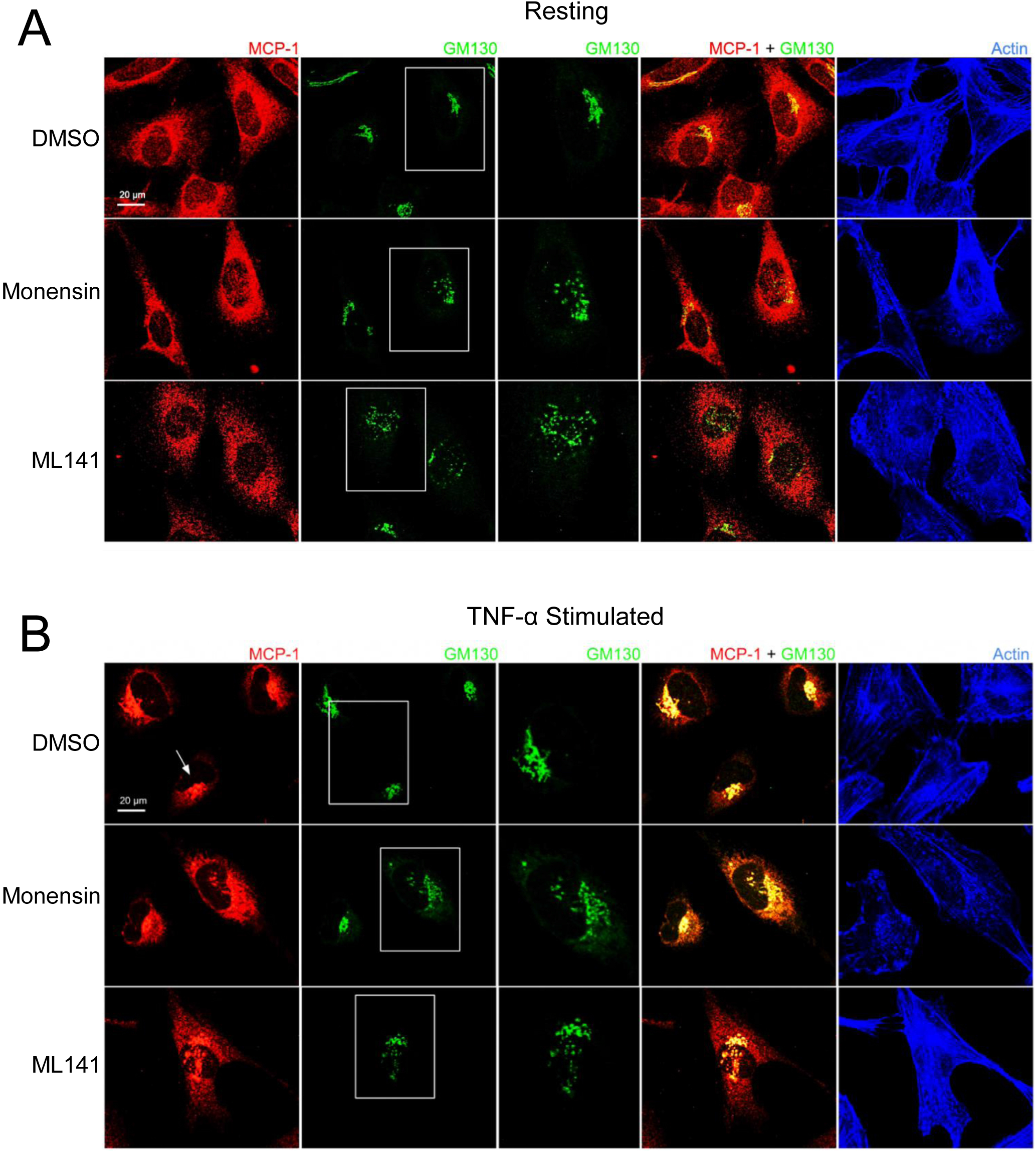
Cdc42 inhibition induces changes in Golgi structure and in MCP-1 trafficking. BEAS-2B cells were pre-treated with 20 μM ML141, 2 μM monensin or vehicle (DMSO) for 1 h. **A)** Cells unstimulated (*resting*) for 4 h. **B)** Cells were stimulated with 10 ng/mL TNF-α for 4 h. Cells were then fixed and stained with MCP-1 and GM130 (Golgi marker) antibodies while actin was labeled with phalloidin-iFluor 405 dye. Monensin did not induced an enrichment of Golgi-localized MCP-1 and ML141 treatment resulted in no change in the MCP-1 staining pattern.

### Depletion of Cdc42 affects cytokine trafficking

Next we used shRNA to deplete Cdc42 in BEAS-2B cells to validate its functional requirement in cytokine production and trafficking. Cdc42 is essential for viability and therefore we selected healthy strains transduced with lentiviral particles encoding shRNAs to three different target sites that showed 40%, 60% and 75% reduction in Cdc42 mRNA (Cdc42 KD strain 1, strain 2, strain 3, respectively) (**Figure 7A**). Generally, Cdc42 knockdown strains showed an increase in IL-8 and IL-1β mRNA levels and a decrease in MCP-1 (**Figure 7B**). The level of Cdc42 knockdown correlated with the increase in IL-8 and IL-1β mRNA levels (**Figure 7B**), which was similar to that observed after treatment with the Cdc42 inhibitor ML141 (*see Figure 3A*). Cdc42 knockdown resulted in decreased MCP-1 levels (**Figure 7B**), which also corroborates the findings of pharmacological inhibition.

**Figure 7.**
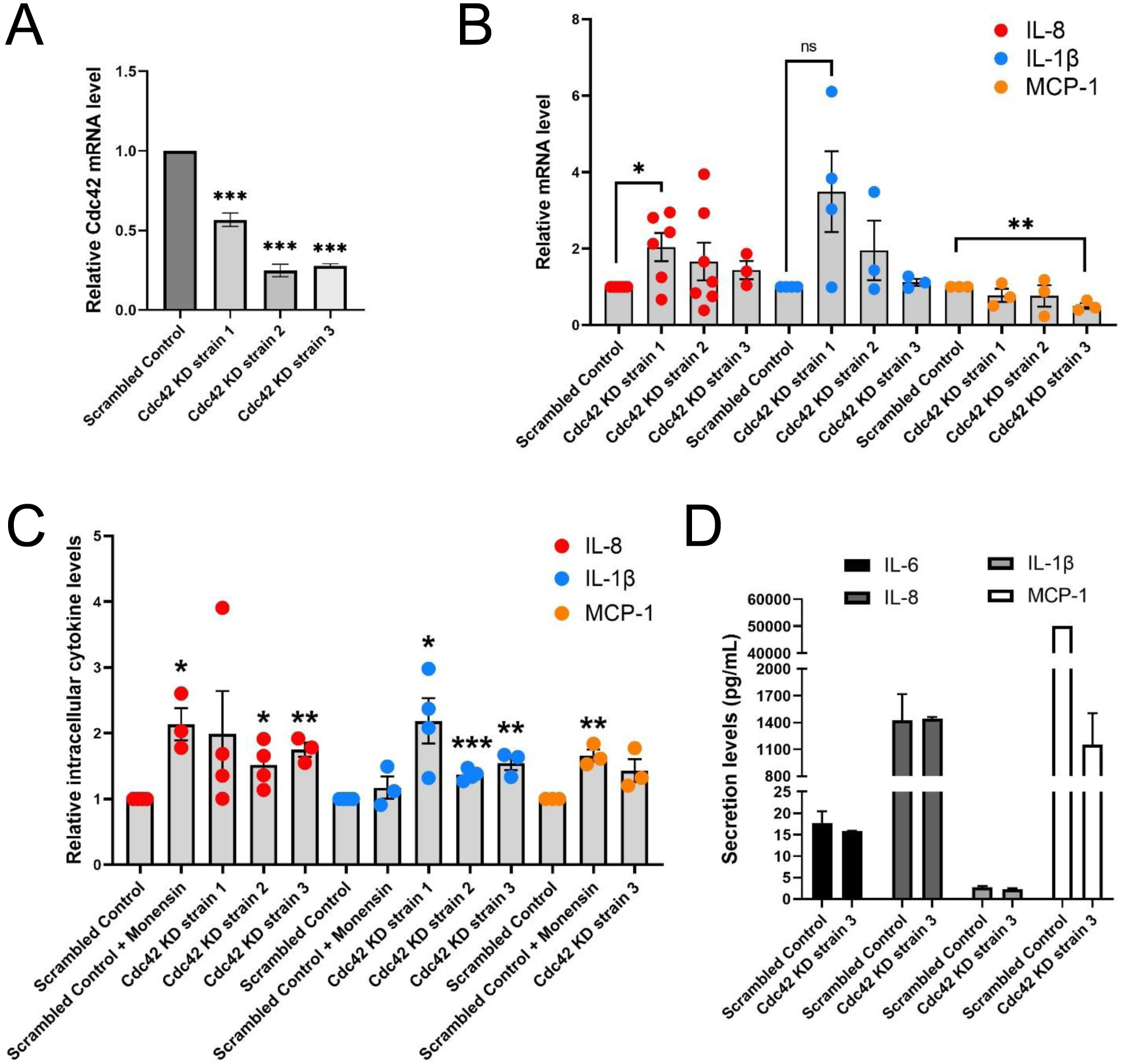
Cdc42 knockdown (KD) affects expression and secretion of a subset of cytokines. **A)** Cdc42 mRNA levels detected by qPCR for three Cdc42 KD strains in BEAS-2B cells. Values represent ΔΔCt fold changes normalized to scrambled control KD. **B)** mRNA levels of various cytokines detected with qPCR in Cdc42 KD strains compared to scrambled control. All cells were serum-starved for 16 h and stimulated with 10 ng/mL TNF-α for 4 h. Values represent ΔΔCt fold changes normalized to scrambled control. **C)** Intracellular levels of cytokine proteins detected with flow cytometry. Serum-starved cells were treated with 2 μM monensin to block secretion or vehicle control (*DMSO*) for 1 h. Cells were then stimulated with 10 ng/mL TNF-α for 4 h. Cells were then trypsinized, fixed for staining and analyzed by flow cytometry. **D)** Secretion levels of IL-6, IL-8, IL-1β and MCP-1 cytokines in cell culture media performed by human cytokine multiplex assay (EveTechnologies™). All cells were serum-starved for 16 h then treated with 10 ng/mL TNF-α for 8 h. Data represents three independent experiments. Bars are the mean ± s.e.m; n=3; *** p < 0.001, ** p < 0.01, * p < 0.05 **, by unpaired two-tailed Student’s t-test. Note, secreted levels of MCP-1 for scrambled control samples were at the upper limit of detection, therefore s.e.m. was not calculated.

To further investigate the effect of Cdc42 silencing on cytokine production, we performed flow cytometry to assess the intracellular levels of IL-8 and IL-1β proteins. Monensin was used as a control to examine the effect of blocking Golgi transport. As expected, levels of IL-8, which traffics through the Golgi (25), increased when monensin treated, while levels of IL-1β, which is constitutively secreted (23), were not affected (**Figure 7C**). Cdc42 knockdown also resulted in an increase in the levels of intracellular cytokines (**Figure 7C**). Surprisingly, we found that MCP-1 intracellular protein levels increase (**Figure 7C**, *yellow dots*), despite the reduction in MCP-1 mRNA levels upon Cdc42 depletion (*see* Figure 7B). This suggests that cytokine accumulation within a cell is due to a block at the trafficking or secretion level, corroborating the findings observed with ML141. Levels of secreted cytokines detected in the extracellular media also generally decreased in Cdc42 KD strains (**Figure 7D** and **Table S2**, *supplementary material*), although we did not detect a reduction in IL-8 which may be due to its increase in transcription. These results support an essential role for Cdc42 in cytokine secretion which appears disrupted when Cdc42 is depleted.

Cdc42 depletion resulted in unique cytokine trafficking disruption as visualized with immunofluorescence microscopy. In resting conditions, IL-8 displayed a characteristic staining pattern with tubules extending towards the cell periphery in the mock-depleted cells (**Figure 8A,** *top panels*). This tubular pattern was greatly reduced by Cdc42 depletion (**Figure 8A**, *bottom panels*). This change was more pronounced with TNF-α treatment, where tubule labelling was eliminated and intense staining was observed at the Golgi with punctate peripheral staining (**Figure 8B**). This suggests that trafficking from the Golgi to the cell periphery is disrupted. MCP-1 showed similar to IL-8. In resting conditions, mock-depleted cells showed a diffuse MCP-1 staining pattern with some tubule label (**Figure 8C***, top panel);* this was eliminated upon Cdc42 depletion (**Figure 8C**, *bottom panel*). Following TNF-α stimulation, MCP-1 displays a characteristic perinuclear staining, which is observed in both mock and Cdc42-depleted cells (**Figure 8D**). No Golgi fragmentation was observed in knockdown cells, as visualized with the Golgi marker GM130 (**Figure 8***, green channel*). This is likely due to cell adaptation because cells with disrupted Golgi cannot survive for prolonged periods of time. This is unlike ML141 treatment, where the effects on the Golgi were acute and cells do not have time to adapt.

**Figure 8.**
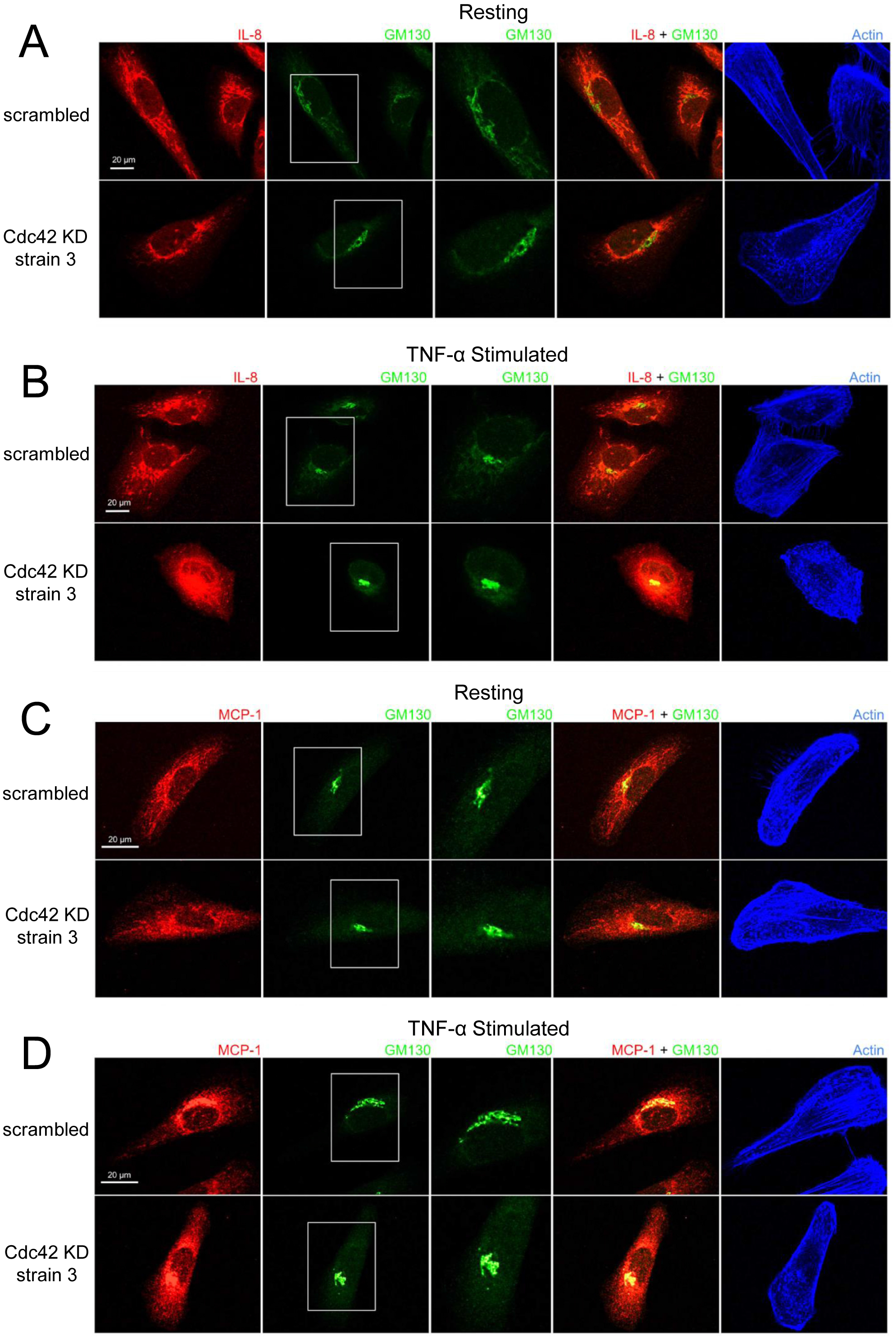
Cdc42 knockdown (KD) induces changes in IL-8 and MCP-1 trafficking. Serum-starved Cdc42 KD and scrambled control BEAS-2B cells were either left unstimulated (*panels A and C*) or stimulated with 10 ng/mL TNF-α for 4 h (*panels B and D*). Cells were fixed and stained with IL-8 or MCP-1 cytokine antibodies, GM130 antibody to label Golgi and phalloidin-iFluor 405 dye to label F-actin. **A and B)** IL-8 staining pattern is tubular in control cells (*scrambled*) but punctate in Cdc42 KD cells, which indicates a significant disruption in trafficking. **C and D)** MCP-1 staining pattern shows Cdc42 KD disrupted MCP-1 trafficking, but had a less significant effect compared to IL-8.

To confirm that the changes observed in IL-8 and MCP-1 trafficking upon Cdc42 silencing are a result of secretory pathway disruptions, we investigated whether the diffuse non-localized staining patterns of IL-1β undergoes similar changes. As expected, IL-1β general cytosolic staining is unaffected and shows no differences in Cdc42-depleted group compared to mock-depleted group (**Figure S2**, *supplementary material*).Another marker, GGA2, that localizes to the *trans*-Golgi Network (TGN) was used to stain the secretory network more broadly (26) highlighting that no significant overlap with IL-1β secretion exists. These effects are consistent with the established mechanisms of trafficking and secretion of IL-1β, highlighting that Cdc42 inhibition may target IL-1β transcription, whereas effects on the secretory pathway do not significantly affect IL-1β.

## DISCUSSION

Several cellular mechanisms protect the airways and lungs from pathogenic substances. Innate immune cells, such as mast cells and macrophage are critical residents of lung tissue that can quickly response to pathogens and infection. The airway epithelium, which primarily provides barrier function, also directly contributes to immune function (27). Studies have shown that airway epithelial cells produce a plethora of cytokines and chemokines with specific profiles in response to specific insults. For example, respiratory viruses trigger interferon production, which in turn upregulate interferon regulated genes to support apoptosis (28, 29), while IL-6 and IL-8 are significantly upregulated in response to viral and bacterial infection to recruit neutrophils for clearance (29–31). However, airway function can be significantly impaired by uncontrolled or allergic inflammation. Consequently, the immune capabilities of the airway epithelium directly contribute to diseases such as asthma and COPD (2). Hence an understanding of how airway epithelial cells regulated immune function may provide therapeutic benefits.

Here we examined how airway epithelial cells regulate pro-inflammatory cytokine production and release. To examine this, we used the human bronchial epithelial cell line, BEAS-2B which we found respond to allergen and inflammatory stimuli through upregulation of cytokines. TNF-α potently upregulated typical epithelial cytokines through NF-κB nuclear translocation, which is known to drive their transcription (32). We found novel roles the Rho GTPase, Cdc42, in the regulation of cytokine transcription and secretion. The effects of pharmacological inhibition of Cdc42 or genetic depletion of Cdc42 found to be somewhat different depending on the cytokine species. While all of these cytokines were transcriptionally upregulated by pro-inflammatory stimuli, inhibition of Cdc42 resulted in upregulation of IL-8, IL-6 and IL-1β, but a dramatic reduction in MCP-1. It seems that only MCP-1 secretion was decreased by Cdc42 inhibition, and there was no effect on IL-8, and IL-6. However, this observation of “no net-change” in IL-6 and IL-8 levels may actually indicate reduced secretion since these cytokines were significantly upregulated by Cdc42 inhibitor treatment (Figure 1B), however their trafficking was disrupted (Figure 5B and 6B) which may counteract the increase in transcription. Another pro-inflammatory cytokine, GM-CSF, that regulates the differentiation and function of immune cells (33) was detected in conditioned media from mock-depleted cells but was not detected in Cdc42-depleted cells (Table S2, *supplementary material*). This suggests that the observed effect of Cdc42 regulation of pro-inflammatory cytokine secretion likely includes a variety of other cytokines or pro-inflammatory cargo proteins, in particular, those using the secretory pathway.

Our results show that Cdc42 has pleiotropic roles in regulating epithelial airway inflammation. In our study, three different cytokines show differential production and trafficking regulation by Cdc42. IL-1β gene expression is positively regulated by Cdc42. Since IL-1β is constitutively secreted, its trafficking does not appear to be affected by Cdc42. On the other hand, IL-8, gene expression is negatively regulated while trafficking is positively regulated by Cdc42. Hence, upon Cdc42 inhibition, it appears as though there is no ‘net effect’ in secretion. This is a result of the antagonism between these two effector functions of Cdc42 that together regulate cytokine release. Thirdly, for the MCP-1 cytokine, gene expression and trafficking are both positively regulated by Cdc42. Therefore, upon Cdc42 inhibition, we observe drastic reductions in secretion as a result of the combined effect of these two effector functions of Cdc42. These three different scenarios suggest that Cdc42 acts as a regulator in different signaling pathways as it acts as a negative transcriptional regulator of some cytokines (such as IL-8, IL-6, IL-1β) and positive regulator of others (such as MCP-1). These results imply that Cdc42 can selectively modulate cytokine production.

Cdc42 also affects trafficking of cytokines, which ultimately impacts their observed secretion. Trafficking regulated by Cdc42 appears to be happening at the level of the Golgi since only MCP-1 and IL-8 trafficking were affected, and not IL-1β. Immunofluorescence microscopy showed prominent disruption to Golgi architecture in response to Cdc42 inhibition (see Figure 5 and 6, GM130 panels after ML141 treatment). Disruption in the staining pattern of the Golgi concur with previous findings showing that pharmacological inhibition of Cdc42 with the small molecule inhibitor, ZCL278, resulted in disruption to Golgi organization in Swiss 3T3 cells (34). Upon treatment with ZCL278, the characteristic perinuclear staining of GM130, a cytoplasmic protein that binds Golgi membranes, is dispersed to both sides of the nucleus. These results imply that Cdc42 may be a highly selectively modulator of cytokine release, highlighting the significance of its therapeutic potential. Indeed, characterization of targets that allow the precise control of the cytokine profile secreted from epithelial cells can prove extremely beneficial in a variety of pathologies. Here, we show that the release of the neutrophil chemoattractant, MCP-1, is attenuated by Cdc42. This is particularly important in COPD where the neutrophil-driven inflammation underlies tissue damage and lung function decline (35). This effect can also be important in lung infections since the cytokine profile can be regulated more selectively to facilitate an antiviral response (mounted by interferons) and directed away from cytokine storms (mounted by pro-inflammatory cytokines).

We report a specific role for Cdc42 in the trafficking of cytokine protein products at the Golgi. A role for Cdc42 in regulating the Golgi trafficking of proteins is expected given the presence of a Golgi pool of Cdc42 in addition to a plasma membrane pool (36). Our immunofluorescence data suggests that Cdc42 is required for proper Golgi architecture. Cytokine trafficking patterns of IL-8 and MCP-1 were disrupted upon Cdc42 inhibition by ML141 or Cdc42 depletion. These cytokines showed transport in post-Golgi tubules like structures that became dispersed punctate after Cdc42 inhibition. This is consistent with findings showing that Cdc42 selectively enhances anterograde transport at the Golgi while inhibiting retrograde transport (37). Specifically, Cdc42 can promote the formation of tubules at the Golgi while inhibiting vesicle formation in the retrograde direction.

Although results from genetic silencing of Cdc42 were by and large similar to that observed with pharmacological inhibition via ML141, not all of the findings were entirely consistent; there was a lack of Golgi fragmentation when Cdc42 was depleted (Figure 8). This is likely due to adaptation that can take place in knock-down cells that are cultured several days compared to the hour of acute treatment with ML141. Cells likely faced high pressure to adapt or compensate for perturbations such as Golgi disruption, increased cytokine production and the reduction of Cdc42 which is an essential protein.

Similarly, Cdc42 KO in fibroblastoid cell models did not show disruptions in Golgi structure where the percentage of cells displaying a compact Golgi stayed the same compared to control (38).

In summary, we report two roles for Cdc42 as it can both negatively and positively affect cytokine gene transcription and concomitantly regulate trafficking and secretion of cytokine protein products. These novel roles for Cdc42 have implications in allergic pathologies, chronic inflammation as well as in infections.

## MATERIALS AND METHODS

### Cell culture and drug treatment

BEAS-2B and HEK-293T cells (ATCC) were grown in DMEM (Sigma) supplemented with 10% heat-inactivated fetal bovine serum (FBS) (Gibco), 10 units/ml penicillin, 100 μg/ml streptomycin and 0.25 μg/ml amphotericin B (Gibco). Cells were grown in a humidified incubator at 37°C and 5% CO_2_. Cells were serum-starved in FBS-free DMEM medium overnight. Prior to stimulation, BEAS-2B cells were pretreated with the drugs, ML141 (Tocris), rhosin (Tocris), EHT1864 (Tocris), monensin (Sigma) and BAY 11-7082 (Cayman) at concentrations indicated in figure legends, or DMSO (vehicle control) for 1 h. Cells were subsequently stimulated with TNF-α at 10 ng/mL or poly(I:C) (NovusBio) at 10 μg/mL or cockroach extract at 20 μg/mL (a gift from Dr. H. Vliagoftis, Alberta Respiratory Centre) for 4 h for microscopy and qPCR analysis, and 8 h for secretion analysis.

### Lentivirus production

Transfer plasmids with pre-designed shRNA against human Cdc42 mRNA were obtained from the Mission^®^ TRC library (Sigma). Specifically, transfer plasmids #34022 (Cdc42 KD strain 1) #34025 (Cdc42 KD strain 2) and #34021 (Cdc42 KD strain 3) were used. The human embryonic kidney cell-line 293T (HEK-293T cells) was used for the production of shRNA lentivirus particles. Briefly, 9 μg of shRNA transfer plasmid, 6 μg of psPAX2 packaging plasmid, 3 μg of pMD2.g VSV-G envelope plasmid and 72 μg polyethylenimine were mixed in 1 ml Opti-MEM, then added dropwise to HEK-293T cells growing in a 10 cm dish at 60% confluency. Lentivirus was harvested 48 h after transfection and filtered by 0.45 μm syringe filter. For the transduction of cells, viral supernatants were added to cells with 10 μg/mL polybrene (hexadimethrine bromide, Sigma) at an MOI of 10 - 15.

### qPCR

Total RNA was isolated using Trizol (Ambion™) according to the manufacturer’s instructions. RNA concentrations were measured using a NanoVue spectrophotometer (GE Life Sciences). Complementary DNA (cDNA) was synthesized from 2 μg RNA and 0.4 μg oligo dT in a volume of 20 μl, using the SuperScript™ II Reverse Transcriptase kit (Invitrogen) according to manufacturer’s protocol. Cytokine and Cdc42 mRNA levels were determined using Mastercycler RealPlex 2 (Eppendorf) thermocycler and SensiFAST™ SYBR No-ROX Kit (Meridion) according to manufacturer’s instructions. GAPDH was used as the internal control reference for the purpose of normalizing mRNA levels. Primer sequences are listed below:

Homo sapiens **IL-8** <u>forward primer</u> (5’ CCAAGGAGTGCTAAAGAACTTAGA 3’) and <u>reverse primer</u> (5’ GTGTGGTCCACTCTCAATCAC 3’).

Homo sapiens **IL-1β** <u>forward primer</u> (5’ CAAAGGCGGCCAGGATATAA 3’) and <u>reverse primer</u> (5’ CTAGGGATTGAGTCCACATTCAG 3’).

Homo sapiens **MCP-1** <u>forward primer</u> (5’ GGCTGAGACTAACCCAGAAAC 3’) and <u>reverse primer</u> (5’ GAATGAAGGTGGCTGCTATGA 3’).

Homo sapiens **IL-6** <u>forward primer</u> (5’ CACTCACCTCTTCAGAACGAAT 3’) and <u>reverse primer</u> (5’ GCTGCTTTCACACATGTTACTC 3’).

Homo sapiens **Cdc42** <u>forward primer</u> (5’ AAAGTG GGTGCCTGAGATAAC 3’) and <u>reverse primer</u> (5’ TGGAGTGATAGGCTTCTGTTT 3’).

Homo sapiens **GAPDH** <u>forward primer</u> (5’ GGTGTGAACCATGAGAAGTATGA 3’) and <u>reverse primer</u> (5’ GAGTCCTTCCACGATACCAAAG 3’).

### Immunofluorescence microscopy

150,000 cells were seeded on cover slips and treated as indicated in figure legends. At the end of stimulation/treatment time, cells were washed into PBS and fixed with 4% paraformaldehyde for 20 min and permeabilized with 0.2% Triton X-100 for 15 min and blocked in 1.5% BSA (Fisher). Cell were labelled with rabbit anti-IL-8 antibody (Biorad Biosciences), mouse anti-IL-1β antibody (Invitrogen), mouse anti-MCP-1 antibody (Abcam), mouse anti-NF-κB p65 antibody (Santa Cruz Biotechnology), mouse anti-GM-130 antibody (BD Biosciences) or rabbit anti-GGA2 antibody (BD Biosciences), and secondary Alexafluor 555 donkey anti-rabbit and Alexafluor 488 donkey anti-mouse antibodies (Invitrogen) and phalloidin-iFluor 405 (Abcam) or DAPI (Sigma). Cover slips were mounted with Mowiol mounting media (Gift from Dr. A. Simmonds, University of Alberta) and imaged with a Zeiss LSM 700 confocal microscope (Zeiss) using a 63x objective (NA 1.4) and images acquired using Zen software (Zeiss). Images were subsequently processed with ImageJ Software (U.S. National Institutes of Health).

### Flow cytometry

BEAS-2B cells were pretreated with ML141, monensin or vehicle control (DMSO) for 1 h. Cells were subsequently stimulated with TNF-α for 4 h, fixed with 4% paraformaldehyde and resuspended in flow cytometry buffer composed of 0.1%Saponin (Sigma), 1% BSA (Fisher), PBS (Sigma). Unstained cells were treated with primary antibodies only and used for gating to identify non-specific signals. Cells were stained with rabbit anti-IL-8 antibody (Biorad Biosciences) and mouse anti-IL-1β antibody (Invitrogen) and Alexafluor 555 donkey anti-rabbit and Alexafluor 488 donkey anti-mouse antibodies (Invitrogen). BD LSRFortessa™ Cell Analyzer was used to detect fluorescence within cells. BD FACSDiva™ Software was used for data acquisition and analysis.

### Cytokine secretion

Levels of secreted cytokines from BEAS-2B cells were determined by analyzing extracellular supernatants. BEAS-2B cells were left resting or stimulated with cockroach extract, poly(I:C) or TNF-α for 8 h. Following stimulation time, media samples were collected and sent for analysis using the Human Cytokine Proinflammatory Focused 15-Plex Discovery Assay^®^ Array (HDF15) (EveTechnologies™). To determine the effect of Cdc42 inhibition or knock-down, BEAS-2B cells were pretreated with ML141 drug or vehicle control (DMSO) for 1 h, or shRNA against Cdc42 5 - 8 days prior to analysis. Cells were subsequently stimulated with TNF-α for 8 h before collecting extracellular media.

### Statistical Analysis

Values reported are the statistical mean +/− standard error of the mean. We used unpaired, two-tail Student’s t-test to determine if the differences between two datasets were statistically significant (p-values). P-values less than 0.05 were deemed significant.

## ACKNOWLEGEMENTS

We would like to thank Drs. A. Simmonds and H. Vliagoftis (University of Alberta) for the gift of reagents. The shRNA from Mission^®^ TRC library were provided by the University of Alberta Faculty of Medicine & Dentistry High Content Analysis Core, RRID:SCR_019182, which receives financial support from the Faculty of Medicine & Dentistry, the Li Ka Shing Institute of Virology, and Canada Foundation for Innovation (CFI) awards to contributing investigators. This work was support by a Discovery Grant from the Natural Sciences and Engineering Research Council of Canada (RGPIN-2019-05466) to G.E. R.S. was supported by a Canada Graduate Scholarship. The funders had no role in study design, data collection and analysis, decision to publish, or preparation of the manuscript.

---

**Supplemental Table 1.** BEAS-2B cytokine secretion levels (pg/mL) after 8 h of stimulation.

|  | Replication 1 |  |  |  |  | Replication 2 |  |  |  |  | Replication 3 |  |  |  |  |
| --- | --- | --- | --- | --- | --- | --- | --- | --- | --- | --- | --- | --- | --- | --- | --- |
|  | Resting | polyI:C | CE | TNF-α | TNF-α + ML141 | Resting | polyI:C | CE | TNF-α | TNF-α + ML141 | Resting | polyI:C | CE | TNF-α | TNF-α + ML141 |
| GM-CSF | N/A | N/A | N/A | 18.2 | N/A | N/A | N/A | N/A | N/A | N/A | N/A | N/A | N/A | 20.7 | 2.19 |
| IFN-γ | N/A | N/A | N/A | N/A | N/A | N/A | N/A | N/A | N/A | N/A | N/A | N/A | N/A | 0.3 | N/A |
| IL-1β | N/A | N/A | N/A | 0.43 | N/A | 1.7 | 1.1 | 0.8 | 2.3 | 2.6 | 1.1 | 1.9 | 1.3 | 2.1 | 1.9 |
| IL-Ra | N/A | N/A | N/A | N/A | N/A | N/A | N/A | N/A | N/A | N/A | 0.5 | 0.6 | 0.6 | 0.7 | 0.6 |
| IL-2 | N/A | N/A | N/A | N/A | N/A | N/A | N/A | N/A | N/A | N/A | N/A | N/A | N/A | 0.2 | 0.2 |
| IL-4 | N/A | N/A | N/A | N/A | N/A | N/A | N/A | N/A | N/A | N/A | N/A | N/A | N/A | N/A | N/A |
| IL-5 | N/A | N/A | N/A | N/A | N/A | N/A | N/A | N/A | N/A | N/A | N/A | N/A | N/A | N/A | N/A |
| IL-6 | 2.6 | 4.6 | 4.3 | 48.2 | 83.6 | 0.3 | 0.3 | 0.4 | 9.9 | 10.5 | 2.4 | 7.5 | 5.3 | 31.3 | 65.6 |
| IL-8 | 41.5 | 70.2 | 56.0 | 693.0 | 599.3 | 5.3 | 6.7 | 2.3 | 85.3 | 79.6 | 68.4 | 135.5 | 103.7 | 1993.1 | 1348.8 |
| IL-10 | N/A | N/A | N/A | N/A | N/A | N/A | N/A | N/A | N/A | N/A | N/A | N/A | N/A | N/A | N/A |
| IL-12p40 | N/A | N/A | N/A | N/A | N/A | N/A | N/A | N/A | 1.3 | 1.6 | 3.4 | 2.3 | 3.7 | 5.5 | 4.4 |
| IL-12p70 | N/A | N/A | N/A | N/A | N/A | N/A | N/A | N/A | N/A | N/A | N/A | N/A | N/A | N/A | N/A |
| IL-13 | N/A | N/A | N/A | 3.57 | N/A | N/A | N/A | N/A | N/A | N/A | N/A | N/A | N/A | 1.31 | N/A |
| MCP-1 | 90.1 | 116.9 | 120.8 | 1625.0 | 255.9 | 7.7 | 8.1 | N/A | 144.8 | 37.5 | 76.4 | 78.1 | 126.5 | 1949.1 | 332.7 |
| TNF-α | N/A | N/A | N/A | 1584.5 | 1211.2 | N/A | N/A | N/A | 1046.5 | 1172.2 | N/A | N/A | N/A | 7875.1 | 7656.5 |

Human cytokine multiplex analysis (HDF15) performed by EveTechnologies™. BEAS-2B cells were stimulated 8 h with 10 µg/mL poly(I:C), 20 µg/mL cockroach extract (CE) or 10 ng/mL TNF-α. ML141 was used at 20 µM and added to cells one hour prior to stimulation. Data represents three independent experiments. N/A – not detected (secretion levels below detection range).

**Figure S1.**
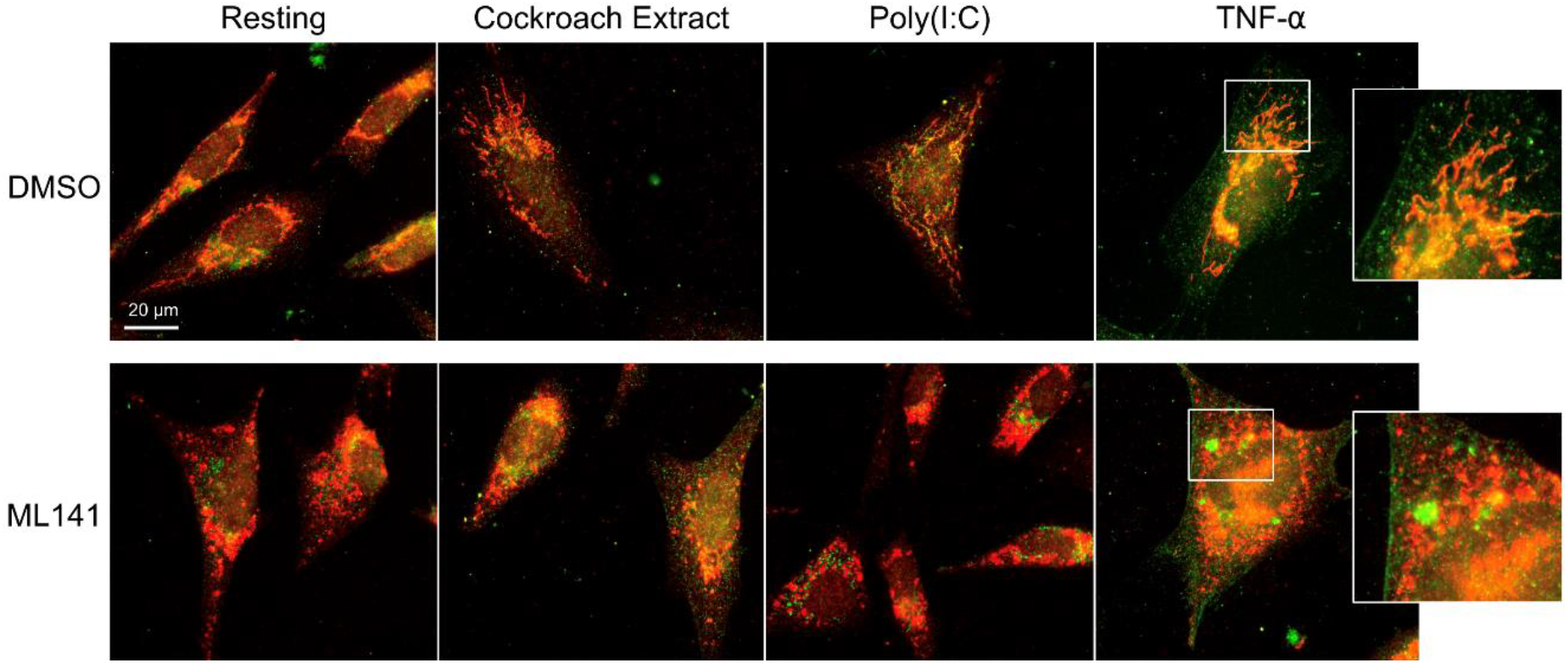
IL-8 and IL-1b show non-overlapping staining patterns and are differentially affected by Cdc42 inhibition. BEAS-2B cells were pretreated with 20 μM ML141 or vehicle (DMSO) for 1 h, then stimulated with 10 ng/mL TNF-α for 4 h. Cells were then fixed and stained with IL-8 (*red*) and IL-1β (*green*) antibodies. Zoomed panels show the two cytokines display different characteristic staining patterns. ML141 affects the IL-8 staining pattern but does not affect IL-1β staining.

**Supplemental Table 2.** Cytokine secretion levels (pg/mL) from BEAS-2B Cdc42 KD and control cells after 8 h of TNF-α stimulation.

|  | Replication 1 |  | Replication 2 |  | Replication 3 |  |
| --- | --- | --- | --- | --- | --- | --- |
|  | ShCtrl | Cdc42-shRNA 3 | ShCtrl | Cdc42-shRNA 3 | ShCtrl | Cdc42-shRNA 3 |
| **GM-CSF** | 6.02 | N/A | 0.50 | N/A | 9.89 | N/A |
| **IFN-γ** | 1.24 | 1.17 | 0.91 | 1.25 | 1.51 | 1.03 |
| **IL-1β** | 2.58 | 2.75 | 2.46 | 1.87 | 3.27 | 2.11 |
| **IL-Ra** | 0.39 | 0.49 | 0.38 | 0.43 | 0.45 | 0.38 |
| **IL-2** | 0.14 | 0.23 | 0.16 | 0.17 | 0.22 | 0.17 |
| **IL-4** | 0.08 | 0.05 | 0.06 | 0.05 | 0.06 | 0.06 |
| **IL-5** | 0.06 | 0.07 | 0.05 | 0.06 | 0.06 | 0.05 |
| **IL-6** | 17.28 | 15.98 | 13.25 | 15.76 | 22.61 | 15.53 |
| **IL-8** | 1509.97 | 1413.76 | 884.23 | 1470.22 | 1883.86 | 1451.87 |
| **IL-10** | 0.38 | 0.35 | 0.30 | 0.32 | 0.44 | 0.34 |
| **IL-12p40** | 4.58 | 5.25 | 2.87 | 4.96 | 5.16 | 4.96 |
| **IL-12p70** | 0.33 | 0.32 | 0.28 | 0.25 | 0.44 | 0.24 |
| **IL-13** | 5.75 | 4.98 | 4.20 | 5.90 | 6.50 | 4.20 |
| **MCP-1** | 50000 | 1856.46 | 50000 | 796.57 | 50000 | 804.62 |
| **TNF-α** | 100000 | 100000 | 100000 | 100000 | 100000 | 100000 |

Human cytokine multiplex analysis (HDF15) performed by EveTechnologies™. BEAS-2B Cdc42 KD or scrambled control cells, were stimulated 8 h with 10 ng/mL TNF-α . Data represents three independent experiments. N/A, not detected (secretion levels below detection range); values in orange color, extrapolated secretion levels based on standard curve (above detection range).

**Figure S2.**
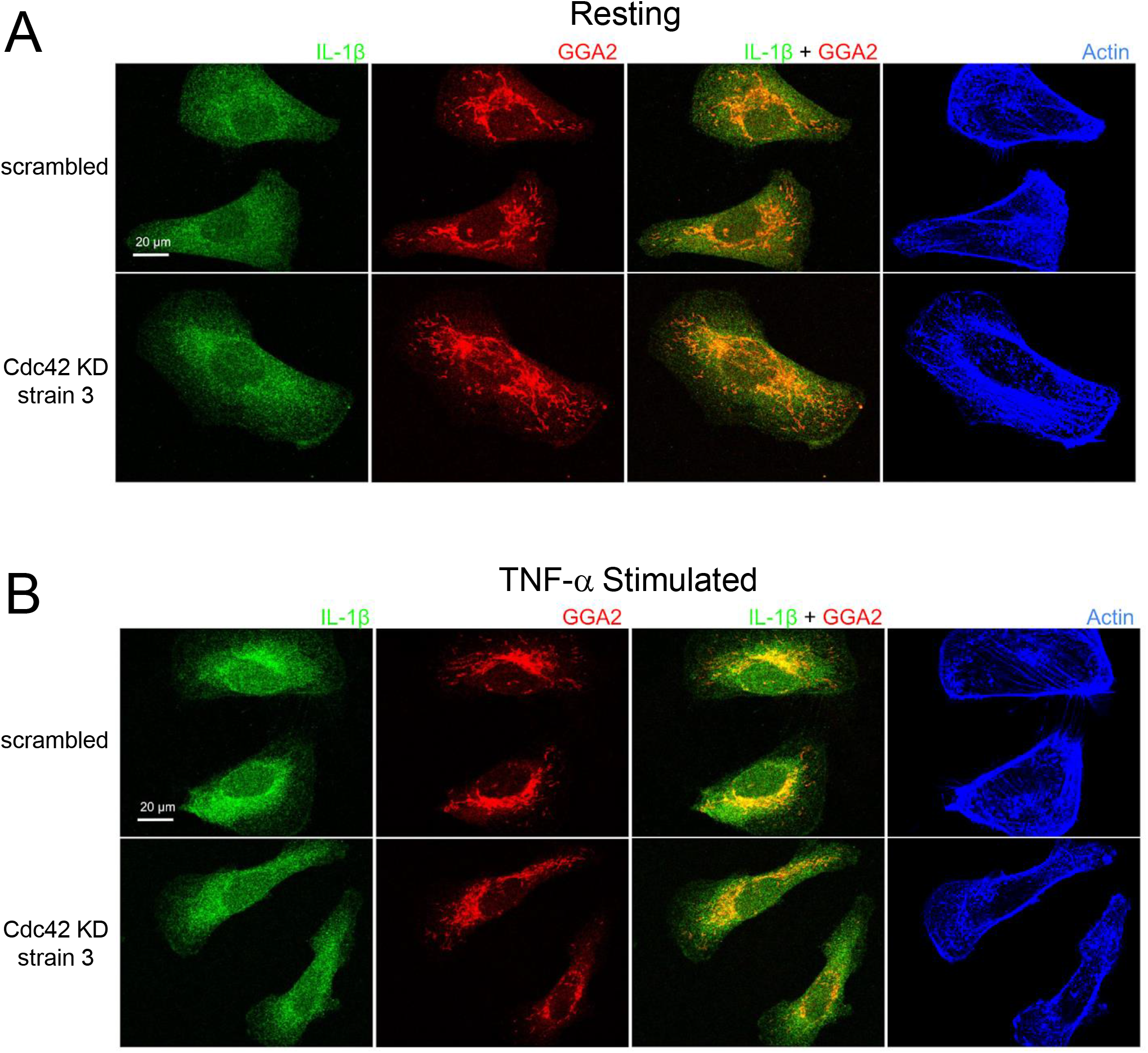
Cdc42-depletion does not affect IL-1β cytosolic trafficking under resting (A) and TNF-α stimulated (B) conditions. BEAS-2B, treated with scrambled control or Cdc42 KD shRNA, were serum-starved overnight, then left unstimulated (*resting*) or stimulated with 10 ng/mL TNF-α for 4 h. Cells were fixed and labeled with antibodies against IL-1 βand the *trans*-Golgi marker GGA3; F-actin was labeled with phalloidin-iFluor405.

